# An ecological synthesis about the effects of floral plantings on crop pollination by bees

**DOI:** 10.1101/2025.09.01.673359

**Authors:** Cristina A. Kita, Isabel Alves-dos-Santos, Michael Hrncir, Marco A. R. Mello

## Abstract

One of the UN’s 17 Sustainable Development Goals is a world free from hunger, with sustainable food production and resilient agricultural practices (SDG 2). However, as we approach the 2030 deadline, global food security, a multidimensional challenge, remains unsolved. With a growing human population, food demand will likely increase, leading to more intensive agricultural practices that put bees and their pollination services at risk. Conserving bees is, thus, crucial to ensuring our food security. A promising solution is cultivating floral plantings alongside crops, which can enhance ecosystem services such as pollination. However, floral plantings may also compete with crops for bees. To optimize their benefits, we need to understand when floral plantings facilitate bee spillover to crops, supporting both bee conservation and agricultural productivity. To address this, we synthesized existing knowledge using a research weaving approach, which combines bibliometric and systematic mapping. Our synthesis suggests that the effects of floral plantings result from a complex interplay of factors, including crop type, bee species, floral planting composition, and environmental conditions. To clarify this interplay, we propose an integrative hypothesis to guide future studies. Finally, our results highlight the need for stronger collaboration among researchers to better understand the role of floral plantings in ecological intensification.

## 1. Introduction

The United Nations (UN) proposed 17 Sustainable Development Goals (SDGs) in 2015, aiming to solve major global problems. Therefore, investing in actions that help us achieve the SDGs is vital to, among other goals, achieving food security, improving nutrition, and promoting sustainable agriculture (SDG 2: Zero Hunger) (United Nations, 2015). That being said, in the coming decades, the human population will continue to grow, increasing our demand for food (Godfray et al., 2010). This increase will potentially intensify even more our agricultural practices (Tscharntke et al., 2012). The dilemma is that agricultural intensification threatens pollinators (Deguines et al., 2014; Freitas et al., 2009). By threatening pollinators, our own food security is put at risk, as our crops largely depend on the service delivered by those animals (Feuerbacher, 2025; IPBES, 2016).

Globally, 74% of animal-pollinated crops are highly dependent on pollinators, and more than 40% of their production is associated with animal pollination, mainly bees (Klein et al., 2007; Siopa et al., 2024). However, pollinators are facing threats worldwide due to several factors, including climate change, pesticides, pathogens, and changes in land use, such as agricultural intensification (IPBES, 2016). Given that intensive crops have expanded rapidly over the past five decades (Aizen and Harder, 2009), reducing vital food and nesting resources for bees, and that bee decline has been reported worldwide (Imperatriz-Fonseca et al., 2012; Zattara and Aizen, 2021), investing in actions to conserve bees and their pollination service in intensive agricultural landscapes has become crucial for ensuring our food and nutrition security through a sustainable food production.

Among these mitigation actions is the use of hedgerows and flower strips adjacent to crops, hereafter named “floral plantings” (von Königslöw et al., 2022). This mitigation action is widely adopted in Europe and North America, where floral plantings are part of transdisciplinary strategies for the conservation of biodiversity in intensively used agricultural landscapes, including bees (Elmiger et al., 2023; Haaland et al., 2011). Thus, the use of floral plantings is highly promising. Hopes are high because floral plantings represent a viable compromise between intensive farming and agroecological systems, providing food, shelter, and nesting sites in resource-scarce agricultural landscapes(Feltham et al., 2015; Häussler et al., 2017). Floral plantings attract bees and can locally enhance their richness, abundance, and diversity, helping to conserve them in agricultural landscapes (Pywell et al., 2011; Venturini et al., 2017).

However, the use of floral plantings adjacent to crops to enhance the pollination service may be tricky. On the one hand, floral plantings can enhance crop pollination as they facilitate the spillover of bees to the crops (hereafter, *exporter hypothesis*) (Blitzer et al., 2012; Kremen et al., 2019). On the other hand, this spillover may not always occur (Nicholson et al., 2019), as floral plantings can concentrate bees for themselves (hereafter, *concentrator hypothesis*), competing with the crops (Bartomeus and Winfree, 2011; Kremen et al., 2019). That is why we need to unveil under which conditions floral plantings can facilitate the spillover of bees to the crops, boosting agricultural productivity in a sustainable way.

Unfortunately, studies on the mechanisms that influence bee dispersal from floral plantings to crops, as well as their drivers and boundary conditions, are still incipient and limited to those two aforementioned competing hypotheses: *concentrator* and *exporter* (Albrecht et al., 2020). Therefore, identifying key factors and integrating those hypotheses into better targeted and logically nested cognitive models (also known as general hypotheses or meta-hypotheses), is an excellent way to deduce finer testable predictions and optimize applied studies. This would not only allow us to fill existing knowledge gaps but also deepen our understanding of the effectiveness of floral plantings in boosting crop pollination. But first, we need to synthesize the knowledge available. One efficient way to do this is through ecological synthesis (*sensu* (Ford and Ishii, 2001)). Syntheses not only allow us to evaluate the state-of-the-art in a field of interest but also produce new knowledge from data and ideas already available, in the form of integrative hypotheses and theories. Therefore, through synthesis, we can more effectively advance knowledge, increasing the return of public investment in science (Kita et al., 2022).

Given the limited knowledge of the effects of floral plantings on crop pollination by bees, and a worrying global scenario of food and nutrition insecurity, our main objective was to help understand and propose directions for optimizing floral plantings to enhance the crop pollination services provided by bees, to indirectly contribute to solving the hunger problem (UN’s SDG 2). While we recognize that hunger is a complex, multidimensional, and transdisciplinary problem, our focus lies specifically on exploring how ecological strategies, particularly improving crop pollination through floral plantings, can contribute to broader efforts to achieve food security within the framework of sustainable agriculture. To fulfill this objective, our goals were to (i) identify, classify, and quantify the evidence available and the main knowledge gaps about the effects of floral plantings on crop pollination by bees, as well as the key mechanisms and factors that influence bee movement from floral plantings to crops; and (ii) assess the development of the field, infer its trends, and identify the key players producing knowledge on the topic, as well as where those studies are being conducted, by mapping co-authorship and institutional networks.

## 2. Methods

To achieve our goals, we used a research weaving approach (Nakagawa et al., 2019). In other words, we conducted a systematic review and, from the articles retrieved, we extracted data to elaborate systematic and bibliometric maps. Thus, we synthesized what has been studied and published to visualize the evidence available and the key players in the field.

### 2.1. Literature search

We searched for peer-reviewed articles on the databases Web of Science (https://www.webofscience.com), Scopus (http://www.scopus.com), and Scielo (https://scielo.org/) including all years up to February 2024. We conducted advanced searches based on title, abstract, and keywords. In these searches, we combined terms related to bees, floral plantings, and agricultural pollination. We searched for terms in English, Portuguese, and Spanish to amplify the literature coverage. Our combination of keywords was: (bee* NOT beetle* OR “wild bee*” OR “native bee*” OR “bee pollinator*” OR “abelha*” OR “abelha* nativa*” OR “abelha* silvestre*” OR “abeja*” OR “abeja* silvestre*”) AND (hedge* OR hedgerow* OR “flower strip*” OR “floral margin*” OR “floral enhancement*” OR “wildflower strip*” OR “field margin*” OR “flower planting*” OR “flowering plant strip*” OR “field edge*” OR “canteiro* de flor*” OR “cerca-viva*” OR “seto*” OR “franja* de flor*” OR “plantación de flor*”) AND (“pollination” OR “crop pollination” OR “pollination service” OR “fruit-set” OR “fruit production” OR “seed*” OR “seed number” OR “seed production” OR “seed-set” OR “polinização” OR “polinização agrícola” OR “serviço de polinização” OR “formação de fruto*” OR “produção de fruto*” OR “semente*”OR “produção de semente*” OR “número de semente*” OR “conjunto de sementes” OR “polinización” OR “polinización agrícola” OR “servicio de polinización” OR “semilla*” OR “número de semilla*” OR “conjunto de semillas” OR “producción de semilla*” OR “formación de fruto*” OR “producción de fruto*”).

### 2.2. Screening

On the selected databases, we identified 1,535 articles. Initially, we imported the complete list into a data frame written in R language (R Core Team, 2024). Using the *litsearchr* package (Grames et al., 2019), we removed duplicates (N = 408). Then, we manually inspected the resulting list (N = 1,070).

Following the screening stages proposed by the Preferred Reporting Items for Systematic Reviews and Meta-Analyses statement (Page et al., 2021) (PRISMA: http://www.prisma-statement.org/), we divided the screening into two stages and pre-established inclusion criteria for each of them. All articles that did not meet those criteria were excluded.

In the first stage, we read the title and abstract of the 1,070 remaining articles. Then, only the articles that met the following criteria were included in the second stage:

1. The study must have been conducted in an agricultural landscape;
2. The study must be related to crop pollination;
3. The study must have investigated the effect of floral plantings adjacent to crops on pollinators or pollination;
4. If the pollinator is identified, it must be a bee;
5. The study must be empirical and conducted under agricultural conditions.

After applying these first inclusion criteria, 200 articles remained in our data set. Besides those articles, during the first stage, we also identified four reviews related to the effect of floral plantings on the bee community. Then, we manually checked the list of articles presented in them (N = 343) and removed duplicates (N = 232). The remaining 111 articles were incorporated into our data set with 200 articles and duplicates were removed again (N = 42). After that, 269 articles were included in the second stage.

In the second stage, we read their methods and results sections. At this stage, only the articles that met the following criteria were included in our review:

1. The study must be empirical and conducted under field conditions;
2. The methodological design must be clearly explained;
3. The study must have reported information that allows us to compare the mean structural metrics of bee communities (e.g., richness and abundance) of a control group with those of a treatment group (e.g., crop with flower strip and crop without flower strip);
4. The crop must have been identified;
5. The crop must depend, to some extent, on bee pollination;
6. The study must have investigated the effect of floral plantings on bee communities within the crops;
7. The bee communities must be local (e.g., nests must not have been introduced into the crop).

After those steps, 25 articles remained in our dataset and were considered suitable for our review (Fig. 1). The full list of articles included in our systematic review is available in Supplementary Material: Articles included in the review.

**Fig. 1.**
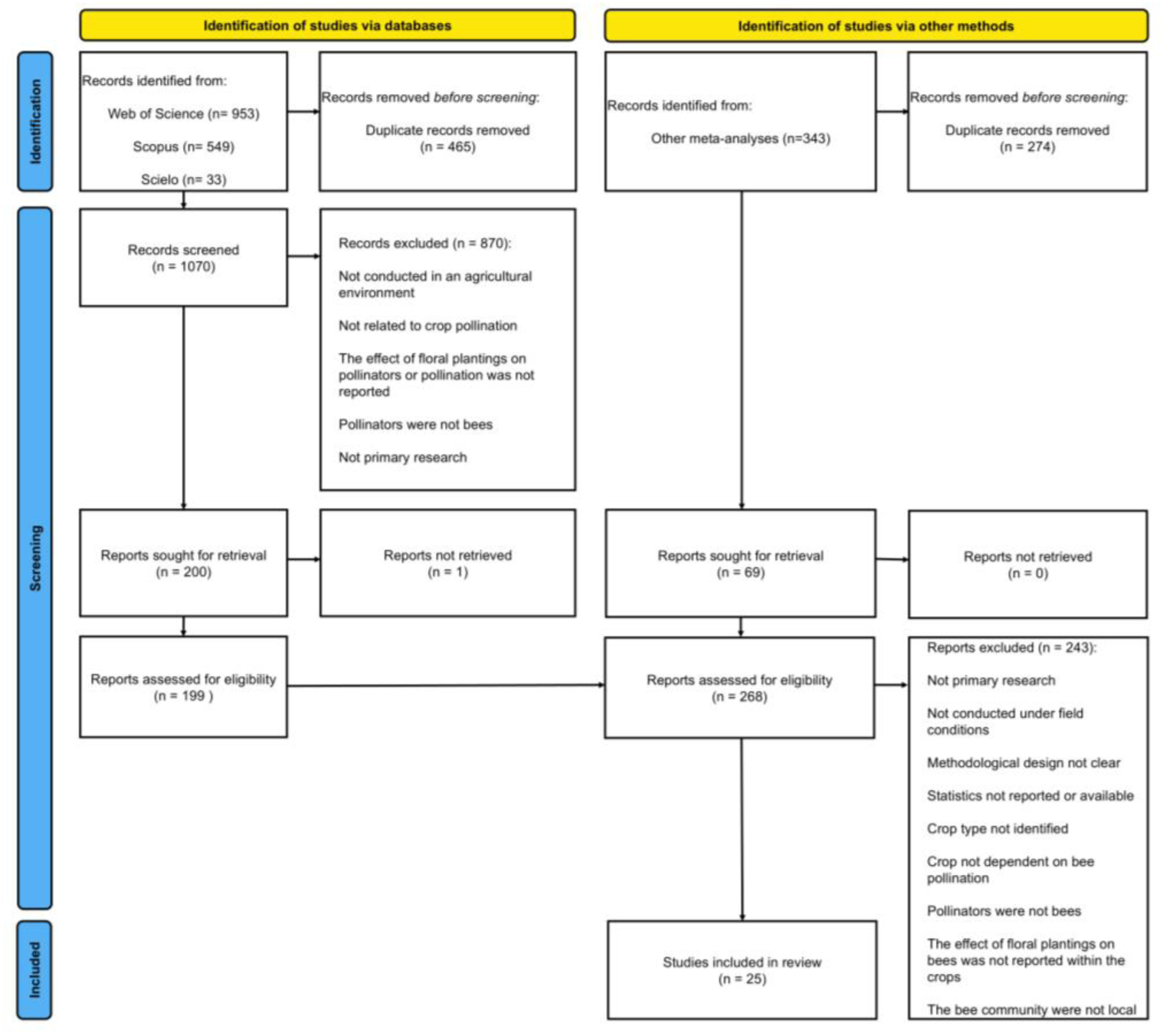
PRISMA flow. Screening flow of our systematic review based on PRISMA.

## 3. Data extraction

Data were manually extracted from article texts and tables. Data extraction was divided into two categories: (i) systematic data and (ii) bibliometric data. In the systematic data extraction, we focused on the content of each publication (e.g., what crops were studied) to construct a content map. In the bibliometric extraction, we focused on the content of the publication itself (i.e., bibliographic and scientometric data) to construct an influence map.

### 3.1. Systematic data extraction

First, we focused on identifying the crops where the studies were conducted, the floral planting composition, and the bee community. We extracted the following data: crop identity, plant species cultivated in the floral plantings (i.e., floral planting composition), and the most abundant bee species collected. We extracted only the most abundant bees because the whole list of bee species collected in each crop was rarely reported whereas the most abundant bee was commonly reported.

Second, we focused on classifying the effects of floral plantings on crop pollination. To do it, we used the number of bees (i.e., abundance), bee species (i.e., richness), and bee visits (i.e., visitation rate) to classify the effects of floral plantings into three categories: (i) exporter, (ii) concentrator, or (iii) neutral. By assessing bee abundance, richness, and visitation rates, we can gain valuable insight into how floral planting and crops may compete with one another for bees, in addition to inferring how bees move between them. This knowledge can shed light on the factors and mechanisms driving bee movement from floral plantings to crops, which might, in turn, influence crop pollination.

Thus, we compared the number of bees, bee species, and bee visitation rate of a control group with those of a treatment group. The crop without floral plantings was considered the control, and the crop with floral plantings was considered the treatment. For instance, if the mean number of bees collected within a crop with floral plantings (treatment) was lower than in a crop without floral plantings (control), we classified the effect of the floral plantings as *concentrator*, meaning that bees moved from the crop to the floral plantings (Fig. 2a).

**Fig. 2.**
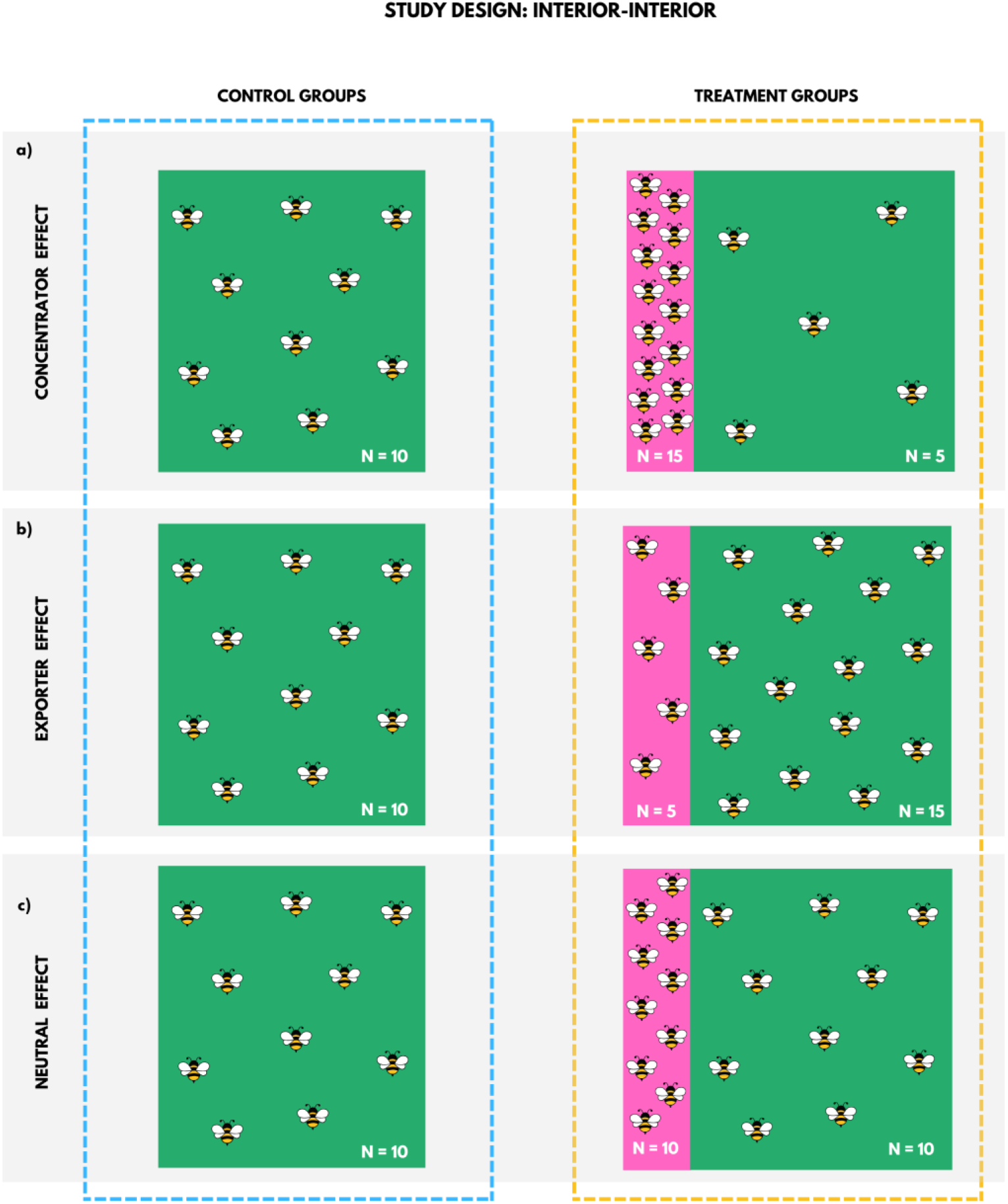
*Interior-interior* study design. The expected effects of floral plantings on crop pollination, using the number of bees collected in the crop as a proxy for bee movement between floral planting and crop, and considering studies with an *interior-interior* design. The green area represents the crop. The pink area represents the floral planting. The bees represent the number of bees collected in each area. The control group is the crop without a floral planting. The treatment is the crop with a floral planting. If the number of bees collected in the crop of the treatment was lower than in the crop of the control, we classified the effect as *concentrator* (a). If the number of bees collected in the crop of the treatment was higher than in the crop of the control, we classified the effect as *exporter* (b). Finally, if there was no difference in the number of bees collected between the crops of the treatment and control, we classified the effect as *neutral* (c).

In other words, we assumed that the floral planting attracted bees from the crop and surroundings, competing for bees with the crop. However, if the mean number of bees collected within a crop with floral plantings was higher than within a crop without floral plantings, we classified the effect as *exporter*, meaning that most bees moved from the floral plantings to the crop (Fig. 2b). In other words, we assumed that the floral plantings attracted bees from the surroundings, which then moved to the crop (bee spillover).

Finally, if there was no difference in bee abundance between treatment and control, we classified the effect as *neutral*, meaning bees were equally attracted by floral plantings and crops (Fig. 2c).

We classified the study design of studies that compared a control group with a treatment group of as *interior-interior,* because the authors compared the number of bees, bee species, and bee visitation rate between two crop interiors. We also considered studies that directly compared the number of bees, bee species, and bee visitation rate between the floral planting and the adjacent crop. We classified this type of study design as *edge-interior*. We also considered this study design because it was possible to infer bee movement between floral plantings and crops.

For example, if the number of bees collected was higher in the floral planting than in the adjacent crop, we classified the effect as *concentrator*, meaning that the floral planting attracted bees from the crop and surroundings, but bees preferred to stay in the floral planting (Fig. 3a). However, if the number of bees was higher in the crop than in the floral planting, we classified the effect as *exporter*, meaning that the floral planting attracted bees from the surroundings, but most bees moved to the crop (bee spillover) (Fig. 3b).

**Fig. 3.**
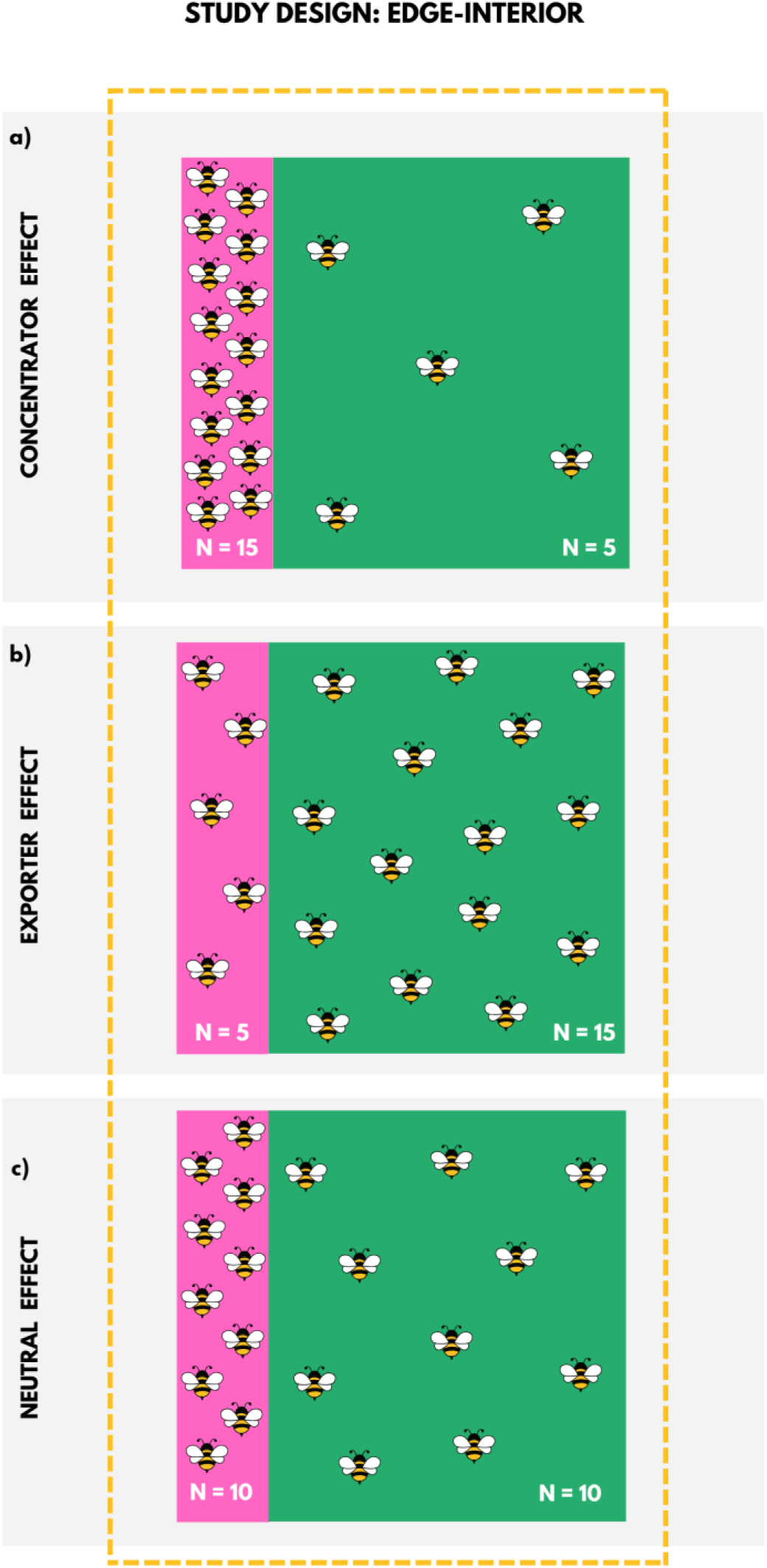
Edge-interior study design. The expected effects of floral plantings on crop pollination using the number of bees collected as a proxy of bee movement between floral planting and crop, and considering studies with an *edge-interior* design. The green area represents the crop. The pink area represents the floral planting. The bees represent the number of bees collected in each area. If the number of bees collected in the floral planting was higher than in the crop, we classified the effect as *concentrator* (a). If the number of bees collected in the floral planting was lower than in the crop, we classified the effect as *exporter* (b). Finally, if there was no difference in the number of bees collected between the floral planting and the crop, we classified the effect as *neutral* (c).

Finally, if there was no difference between the floral planting and the crop, we classified the effect as *neutral*, meaning that the bees did not show any preference for the floral planting or the crop (Fig. 3c). Therefore, we extracted the mean number of bees collected (i.e., abundance), number of bee species collected (i.e., richness), and number of bee visits to the crop (i.e., visitation rate) of the treatment and control groups.

With this information in hand, we classified the resulting effect. We noticed that bees were commonly classified into three main groups: honeybees, wild bees, and bumblebees. As those groups differ from one another in foraging behavior, social structure (sociality), and nutritional preferences (Barraud et al., 2022), we also classified the floral planting effect by bee groups. We expected to detect relationships between crop identity, bee species, and floral planting composition with the effect of floral planting observed.

Finally, we focused on identifying, qualifying, and quantifying other factors related to crop management and features, and landscape configuration and composition of each study site. To do this, we extracted the following data: pesticide use, crop area, percentage of vegetation surrounding the crops, and study sites (i.e., country, state, or crop site in the same state). With this information in hand, we expected to identify relationships between these factors other and the effect of floral planting observed.

### 3.2. Bibliometric map extraction

First, we focused on visualizing the development of the field to identify its trends. We manually extracted the following data: journal of publication, year of publication, and the keywords listed in the studies. To extract the number of citations each study received, we used the Crossref website (https://www.crossref.org).

Second, we mapped co-authorship and institutions to identify the main researchers producing knowledge on the effect of floral plantings as well as their affiliations. We extracted the following data: names of all authors of each study and respective affiliations.

With this information, we expected to identify some biases and collaborations among authors interested in the same topic. All systematic and bibliometric data extracted are available in Supplementary Material: Processed data.

## 4. Data analysis

We made the figures using a combination of R packages to visually analyze the data extracted. Both the systematic and bibliometric maps were drawn by combining plots built separately. All processed data and code, as well as tutorials for facilitating reproducibility, are available in Supplementary Material: Files.

### 4.1. Systematic map

For crop type and floral planting composition, we used the packages *ggplot2* (Wickham, 2016) and *dplyr* (Wickham et al., 2014) to build bar and donut plots. For bee species, we used not only ^3132^the same two packages but also *vdc* (Meyer et al., 2024) to build a pie chart. Finally, for the effects of floral plantings, we combined data related to study site, crop type, floral planting type, and bee group to build a diagram in Canva (https://www.canva.com/).

### 4.2. Bibliometric map

For journal and year of publication, we used *ggplot2* (Wickham, 2016) and *dplyr* (Wickham et al., 2014) to build bar and line chart plots, respectively. For author affiliation, we built a lollipop plot using the same packages. To visualize keywords, we built a word cloud using the packages *dplyr* (Wickham et al., 2014), *reader* (Cooper, 2013), *wordcloud* (Fellows, 2011), and *RcolorBrewer* (Neuwirth, 2022). Finally, for co-authorship, we built a network using the names of authors as nodes and their collaborations as links, using the packages *igraph* (Csárdi et al., 2024), *ggraph (Pedersen, 2024a)*, *ggplot2* (Wickham, 2016), *tibble* (Müller and Wickham, 2023), *dplyr* (Wickham et al., 2014), *ggforce* (Pedersen, 2024b), *randomcolorR* (Ammar, 2016), and *ggrepel* (Slowikowski et al., 2024).

## 5. Results

### 5.1. Systematic mapping

#### 5.1.1. Crops, bees, and floral plantings

In the 25 studies included in our review, we identified 15 crop types: almond, apple, avocado, blueberry, cherry, courgette, field mustard, mango, melon, oilseed rape, red clover, strawberry, sunflower, tomato, and watermelon. The most frequent was blueberry, representing 20% of all studies (Fig. 4a).

**Fig. 4.**
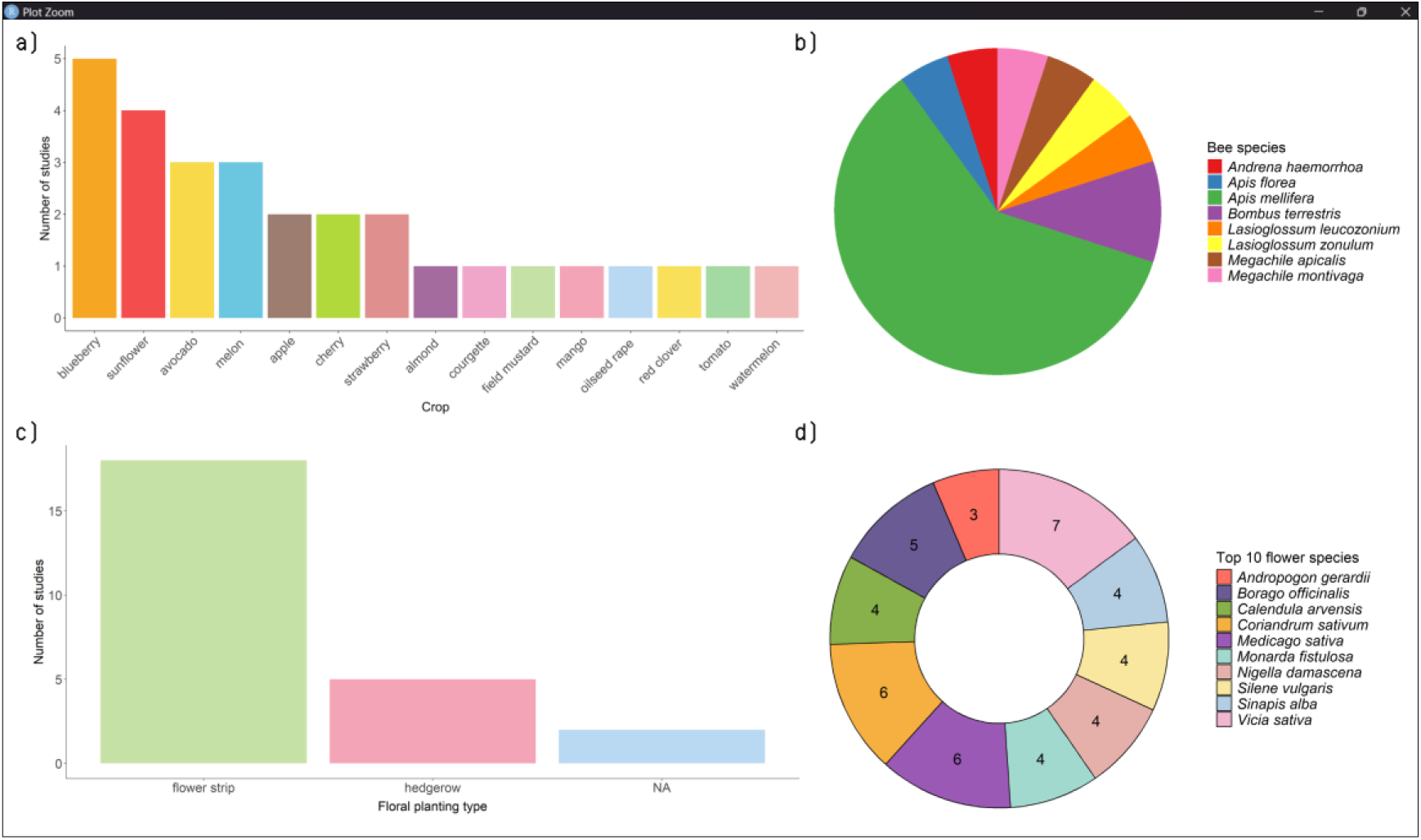
Systematic map. Data extracted from the 25 articles retrieved in our systematic review of the effects of floral plantings on crop pollination by bees. Blueberry was the most frequently studied crop (N = 5) (a); the honeybee (*Apis mellifera*) was the most abundant bee species collected in the crops (b); the most common floral planting type was flower strip (N = 18) (c); and common vetch (*Vicia sativa*) was cultivated in seven of the floral plantings (d).

The bee species collected varied between crop types and study sites. Overall, the most abundant bee species was the managed honeybee *Apis mellifera* (Fig. 4b). This species was the most abundant in almond, apple, avocado, blueberry, courgette, field mustard, melon, oilseed rape, strawberry, and tomato crops. Among wild bees, all of them were solitary bees, except for bumblebees. Out of the 25 studies, four reported only the bee genus of the most abundant species collected (*Bombus* spp. and *Lasioglossum* spp.), and 11 did not report the most abundant bee species.

Floral plantings varied in type (i.e., hedgerow or flower strip). Five studies assessed hedgerows and 18 assessed flower strips (Fig. 4c). Only three studies did not identify the type of floral planting assessed; they reported the presence of vegetation strip, but did not provide further details. Floral planting composition varied both between crop types and within the same crop. Overall, we identified 204 plant species. Eight plants were not identified to the species (*Brassica* sp., *Ceanothus* sp., *Fumaria* sp., *Hyacinthoides* sp., *Lamium* sp., *Malus* sp., *Medicago* sp., and *Papaver* sp.). The three most common plant species cultivated in floral plantings were: common vetch (*Vicia sativa*, Fabaceae), alfalfa (*Medicago sativa*, Fabaceae), and cilantro (*Coriandrum sativium*, Apiaceae) (Fig. 4d).

These species were cultivated with other plants in floral plantings adjacent to the following crops: apple, avocado, melon, strawberry, and sunflower. Out of 25 studies, three did not report the plant species cultivated in the floral plantings.

#### 5.1.2. The effects of floral plantings on crop pollination

The effects of floral plantings on crop pollination varied both across and within studies. We found that 13 studies identified multiple effects within the same study (e.g., combinations of concentrator, exporter, and neutral effects), while the remaining 12 studies reported only a single effect (either concentrator, exporter, or neutral).

We noticed that depending on the bee group analyzed, the effects of floral plantings differed, which suggests that their effects depend not only on the resources available in the floral planting or crop but also on bee group preference and behavior. However, bee group preference might not be the only factor that influences floral planting effects. We observed that for studies conducted in consecutive years (N = 3), the effect of floral plantings varied. In these studies, the crop type, bee group, and floral planting composition did not change between years. This inconsistency suggests that other factors such as environmental conditions may also play a role.

We did not find a consistent pattern of effects that could be explained by a single factor alone (crop type, floral planting composition, or bee group). Instead, the observed effects appear to result from an interplay of factors (crop type, floral planting composition, and bee group) (Fig. 5).

**Fig. 5.**
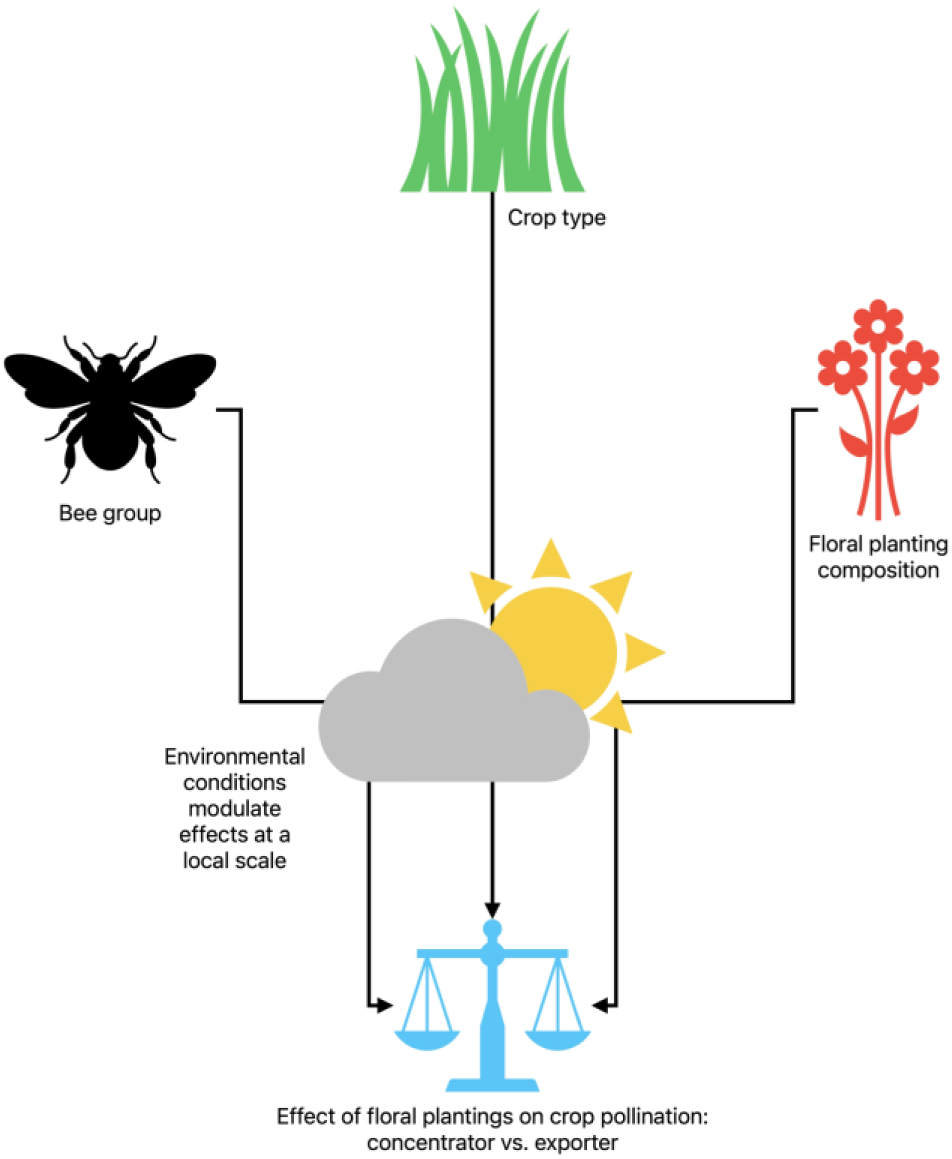
The interplay of factors. Main factors that seem to determine the effects of floral plantings (i.e., concentrator or exporter), as observed in our review.

#### 5.1.3. Other factors and knowledge gaps

The effect of floral plantings did not vary with pesticide use, crop area or proportion of vegetation cover surrounding the crop. However, the number of studies that reported this kind of information was small, so we cannot rule out their potential influence. Below, we highlight the main knowledge gaps detected.

Regarding pesticide use, only 12 studies reported this information. The mean crop areas reported ranged from 0.03 to 11.2 ha, but only 14 studies reported this information. Information on the proportion of natural vegetation surrounding the crops was also scarce. Only four studies reported quantitative information on the proportion of natural or semi-natural vegetation, which ranged from 1 to 40%. Two of these studies mentioned different proportions of semi-natural vegetation surrounding the control and treatment crops, in apple and blueberry crops. Four studies reported information on other crops surrounding the target crop but only described the landscape as a mosaic of large-scale plantations. The other studies did not report any information about landscape features.

Besides the missing information on landscape features, we also identified knowledge gaps related to the countries where the studies were conducted. Overall, the studies were conducted in 12 countries. Most studies were conducted in the U.S.A. (N = 8) and in Spain (N = 5), which shows a bias towards North America and Europe. In the U.S.A., the most common study site was Michigan. In Spain, the most common study sites were Burgos and Cuenca. The other countries reported were Canada, Chile, Ireland, Kenya, Morocco, Netherlands, Sweden, Switzerland, Thailand, and United Kingdom (Supplementary Material: Supplementary text – Fig. 1). Thus, there is a lack of information about the effects of floral plantings in most of the world, mainly South America, Africa, and Asia, which highlights the need for more studies in these continents.

### 5.2. Bibliometric mapping

The most frequent journal where the studies were published was Agriculture, Ecosystem and Environment (N = 4) (Supplementary Material: Supplementary text – Fig. 2). Interestingly, the studies published in this journal support the *concentrator hypothesis*. The years of publication ranged from 2011 to 2023 (Fig. 6a).

**Fig. 6.**
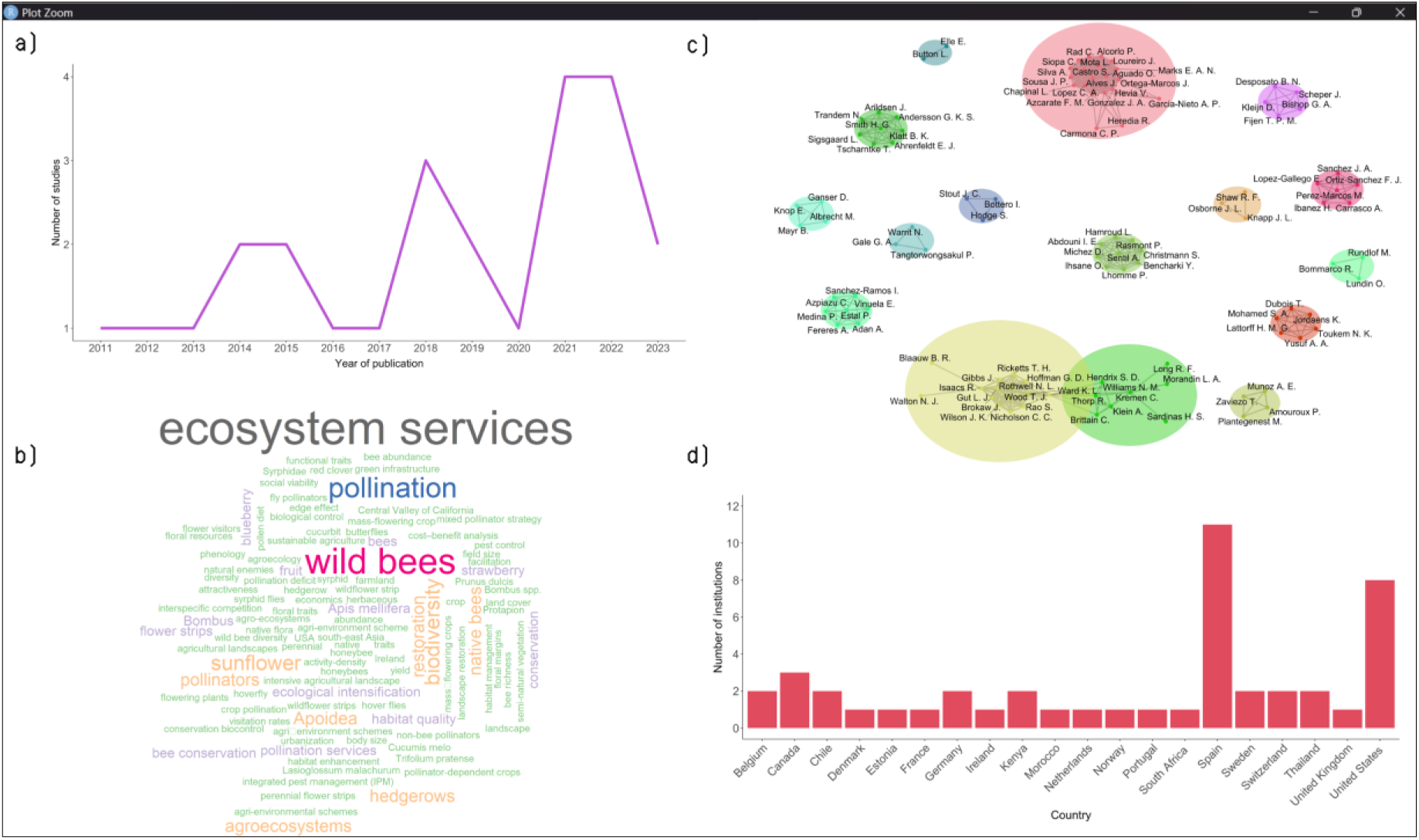
Bibliometric map. Data extracted from the 25 articles retrieved in our systematic review about the effects of floral plantings on crop pollination by bees. The number of studies published per year (a). Keywords listed in the articles retrieved: keyword size represents the frequency of use (b). Co-authorship network: nodes represent authors and links represent collaboration (c). Countries where author’s institutions are located (d).

The number of citations ranged from 1 to 430. The most cited study was Blaauw & Isaacs (2014) with 430 citations, followed by Morandin & Kremen (2013) with 270 citations. Both studies corroborate the *exporter hypothesis* for wild bees. Thus, there may be a bias in the literature towards citing a positive effect of floral plantings on crop pollination by bees, in other words, the exporter effect.

In total, 110 different keywords were used. Contradictorily, the words “flower strip” and “hedgerow” were not the most frequently used. Words related to the *concentrator* and *exporter hypotheses* such as “facilitation”, “spillover”, “competition”, “concentration”, and “bee movement” were not identified. The only word related to these hypotheses was “facilitation” which was used only in one study. The most common keywords used were “ecosystem service” (N = 11), “wild bees” (N = 8), and “pollination” (N = 6) (Fig. 6b).

The author collaboration network was highly disconnected, with many isolated components and a few modules. It means that the key players producing knowledge about floral plantings are not yet intensively collaborating with one another (Fig. 6c). Regarding the affiliated institutions, most authors work in Europe and North America. Institutions from South America, Africa, and Asia were missing, except for two institutions in Chile, two in Kenya, one in Morocco, and two in Thailand (Fig. 6d).

## 6. Discussion

We synthesized knowledge about the effects of floral plantings on crop pollination by bees, aiming to contribute to solving the hunger problem (UN’s SDG 2), a multidimensional puzzle that also requires ecological pieces. Using a research weaving approach, we made systematic and bibliometric maps to help guide future basic and applied studies. Our findings point to promising research avenues.

First, our systematic map suggests that the effects of floral plantings emerge from a complex interplay of factors, mainly crop type, floral planting composition, bee group, and environmental conditions. It should be noted that environmental conditions seem to play a crucial role in regulating the effects of floral plantings at a local scale, but further studies are needed to clarify their influence. Second, our bibliometric map revealed a highly disconnected co-authorship network, which evidences the urgent need for improving collaboration and communication in the field.

Problems in communication are worrisome not only among data producers but also between them and data users, as the lack of standardization in the information reported seriously restrained the scope of our assessment and will probably also hinder other attempts at synthesis (Kita et al., 2022). Therefore, data reporting needs improvement to enhance knowledge exchange and integration between producers, users, and stakeholders (see Kita et al. (2022) (Kita et al., 2022) for guidelines). This way, we can improve our understanding and direct our efforts towards optimizing the use of floral plantings to ecologically intensify agriculture and contribute to achieving SDG 2. However, even with the few data available, we can point to knowledge gaps and research avenues.

### (1) Floral plantings vs. crops

We observed that the effects of floral plantings on bees vary between study sites. For example, two studies focused on melon crops, one in Morocco and the other in Spain, corroborated the *exporter* and *concentrator* hypotheses, respectively. Both studies assessed the effects of floral plantings adjacent to melon crops on wild bee abundance, with the main difference between them being the plant species used in the floral plantings. Probably, the key to this difference is the resources provided by different plants to bees in floral plantings compared to crops. Considering that bees optimize the balance between the energy acquired and expended during foraging (Seeley, 1994), both crops and floral plantings offer resources to bees (Bänsch et al., 2020; Rutschmann et al., 2023), and bee foraging behavior is influenced by nectar and pollen quantity and quality (Vaudo et al., 2015), the effect of floral plantings most likely depends on where resource reward is energetically higher: crops or floral plantings. If it is true, in Morocco, the resource reward offered by the melon crops should be higher than by floral plantings (exporter effect). It might be the other way around in Spain (concentrator effect). Thus, future studies could focus on understanding how resources available in floral plantings and crops can influence bee movement between these areas, for example, by quantifying the resource ratio between them.

### (2) The bee group

Another factor that could strongly influence the effect of floral plantings is bee behavior. We observed that different studies showed different effects of floral plantings depending on the bee group analyzed (i.e., honeybees, wild bees, and bumblebees). It suggests that the effect of floral plantings may depend not only on where the resources (e.g., pollen and nectar) are more rewarding (crops or floral planting) but also on bee foraging behavior. For example, results obtained for courgette crops in the same site in the U.K., with the same floral planting composition, point out that the effect of floral plantings varies with bee group. For honeybees and bumblebees that exhibit recruitment behavior (Alves et al., 2023), the study corroborated the *exporter hypothesis*, while for wild bees (only solitary species) the study corroborated the *concentrator hypothesis*. The same difference between bee groups was observed for strawberry crops in Switzerland, field mustard crops in the United States, and oilseed rape crops in Ireland. Thus, considering that different bee groups have different sociality, which affects foraging behavior (Brunet et al., 2023; Grüter and Hayes, 2022), and resource preference (Leonhardt and Blüthgen, 2012), the effects of floral plantings may also depend on the match between the resources available and bee group.

Morphological traits of bees, such as body size, could also influence the effect of floral plantings. Considering that different bee species differ in body size (Chole et al., 2019), body size limits bee foraging distance (Greenleaf et al., 2007), and the distance between floral plantings and crops can vary as well as the area of each crop, the effect of floral plantings may also depend on the bee species present in the local bee community. Thus, future studies could investigate how different bee species could be strategically used to enhance crop productivity, by considering the interplay between floral planting and crop areas and bee foraging behaviors.

### (3) Environmental conditions

Different floral planting effects could also result from differences in environmental conditions between sites, such as landscape configuration and composition (e.g., other crop types available in the surroundings) (Bottero et al., 2023), which may influence bee movement between crops and floral plantings. Thus, studies conducted in the same crop type with the same floral planting composition, but located in different sites, could corroborate different hypotheses. Therefore, future studies should consider the landscape surrounding the crops and floral plantings. To make matters even more complicated, though, the interplay of factors seems to be a little more complex.

In two studies conducted in blueberry crops in two different sites of Michigan, U.S.A., with the same floral planting composition, the floral planting effect was neutral. However, in another study, also in a blueberry crop, in another site in Michigan, but with different floral planting composition, the effect was exporter. These results lead to raising three hypotheses. First, the environmental conditions of the two studies that supported the neutral effect may be similar to one another, whereas the environmental conditions of the study that supported the exporter effect could be significantly different. Second, at the level of species interactions on a small spatial scale, the variation in the effects may stem mainly from differences in floral planting composition. Third, both environmental conditions and floral planting composition may synergistically influence the observed effect. Thus, future studies could use these hypotheses as a framework to clarify the influence of environmental conditions combined with floral planting composition.

### (4) Knowledge gaps

Unfortunately, limitations in data variety and quantity reported in the literature hinder further assessments. Those limitations show that information reporting about data collection methods, raw data, processed data, and model results must be improved as it is key to advance knowledge not only on the effects of floral plantings but also different types of species interactions (Kita et al., 2022). For example, information on landscape features such as landscape configuration and composition could help assess how environmental conditions can influence the effects of floral plantings since landscape features (patterns) can influence ecosystem services such as crop pollination (Viana et al., 2012). Thus, reporting information, for instance, crop and floral planting areas, percentage of natural vegetation surrounding the crops, types of crops in the vicinity of the assessed crop, and information on land management (e.g., pesticide use), is of paramount importance to understanding the factors behind the effects of floral plantings.

Information from other continents besides Europe and North America is also crucial to fill knowledge gaps, as floral plantings might also represent a viable solution for many other countries, particularly in the tropics, given that most studies in our review were conducted in the Temperate Zone. Tropical studies are crucial for both bee conservation and global economic growth, as the region supports a vast diversity of bee species, especially the Neotropics (Freitas et al., 2009), and is a major producer of key *commodities* such as coffee and cocoa (FAO, 2025).

Conducting experiments in different periods is also important, as the same experiment, focused on the same bee group, under the same environmental conditions, but in different years (N = 3), can lead to observing different effects. We observed that, in a study conducted in sunflower crops, in the first year, neither the concentrator nor the exporter hypotheses were corroborated. However, in the second year, the same study conducted under the same conditions corroborated the *exporter hypothesis*. The same incongruence between consecutive years was also observed for melon and red clover crops. Thus, future studies are needed to clarify these conflicting results.

### (5) A new perspective

Considering that (i) the empirical evidence synthesized here seems to corroborate both the *exporter* and *concentrator hypotheses*; (ii) bee movement (dispersal or commuting) from floral plantings to crops is what differentiates them; (iii) and intensive crops are generally hostile to bees due to their low resource diversity (Kremen et al., 2002), while floral plantings attract them (Vaudo et al., 2015), we propose that these hypotheses may not be mutually exclusive. Instead, they could represent sequential stages of bee movement across the agricultural landscape. Thus, the concentrator hypothesis would represent the stage of bee arrival at floral plantings (concentrator stage), while the exporter hypothesis would represent the stage of bee movement from floral plantings to crops (exporter stage). In other words, the concentrator and exporter effects might be stages of the same phenomenon. Thus, the concentrator hypothesis would represent the stage of bee arrival at floral plantings (concentrator stage), while the exporter hypothesis would represent the stage of bee movement from floral plantings to crops (exporter stage). In other words, the concentrator and exporter effects might be stages of the same phenomenon. Furthermore, we must consider that crops often represent a temporary resource source during flowering. Floral plantings, on the other hand, can sustain bees for a long time. Consequently, there is probably a modular structure in the pollination network formed in farms, which involves crops, floral plantings, natural habitat remnants, and other environments available in the landscape.

In addition, considering that (i) the effect of floral plantings seems to emerge from a complex interplay of factors; (ii) the transition from the concentrator to the exporter stage seems to represent an emergent phenomenon (Johnson, 2002) as observed in other complex systems (Dakos et al., 2023); and (iii) thresholds or tipping points are commonly observed in phenomena involving emergence (Dakos et al., 2023; Scheffer et al., 2009), we propose a new perspective. We hypothesize that, for each factor, there must be a threshold at which bees migrate from floral plantings to crops, taking the system from the concentrator to the exporter stage. This threshold is probably related to resource dissimilarity between habitats and seems to be key to understanding the spatially-explicit modularity that potentially occurs in pollination networks at farms, so it might be better understood within the framework provided by the Integrative Hypothesis of Specialization (Mello and Dormann, 2025; Pinheiro et al., 2019). Maybe, in the end, we need to take a step back to see the bigger picture.

## 7. Conclusion

The effects of floral plantings on crop pollination seem to result from a complex interplay between key factors, mainly crop type, plant species composition of the floral planting, bee community composition, and environmental conditions. However, much more information is needed to assess those factors and the mechanism that articulates them. First, we need future studies to report information in a more standardized way to facilitate synthesis. Second, we need not only further experimental studies on the phenomenon but also studies that conceptually integrate the exporter and concentrator hypotheses. After all, for the sake of our own food security, we need to work hard on the ecological intensification of agriculture to find viable compromises between production and conservation.

## Acknowledgments

We are deeply grateful to the authors of all primary studies included in our systematic review, whose empirical work made our synthesis possible. Special thanks go to Astrid Kleinert and Renata Muylaert for their invaluable insight and advice, which helped us see the bigger picture and put CAK’s Ph.D. project in perspective. We also thank Tereza Giannini, Paula Prist, and Silvana Buzato for their insightful suggestions during CAK’s Qualifying Exam. Last but not least, we thank the Stack Overflow community (https://stackoverflow.com/), where we solve most of our coding dilemmas.

## Supplementary Material

All supplements mentioned in the text, as well as data and code to reproduce all graphical and numerical analyses, are available on GitHub: https://github.com/CKita/BeeFlow.git. The data and code to reproduce the systematic and bibliometric maps are organized into separate folders. The supplementary text and processed data are available as standalone files.

